# Data-driven filtering for denoising of TCRpMHC single-cell data: a benchmark

**DOI:** 10.1101/2023.02.01.526310

**Authors:** Helle Rus Povlsen, Alessandro Montemurro, Leon Eyrich Jessen, Morten Nielsen

## Abstract

Pairing of the T cell receptor (TCR) with its cognate peptide-MHC (pMHC) is a cornerstone in T cell-mediated immunity. Recently, single-cell sequencing coupled with DNA-barcoded MHC multimer staining has enabled high-throughput studies of T cell specificities. However, the immense variability of TCR-pMHC interactions combined with the relatively low signal-to-noise ratio in the data generated using current technologies are complicating these studies. Several approaches have been proposed for denoising single-cell TCR-pMHC specificity data. Here, we present a benchmark evaluating two such denoising methods, ICON and ITRAP. We applied and evaluated the methods on publicly available immune profiling data provided by 10x Genomics. We find that both methods identified approximately 75% of the raw data as noise. We analyzed both internal metrics developed for the purpose and performance on independent data using machine learning methods trained on the raw and denoised 10x data. We find an increased signal-to-noise ratio comparing the denoised to the raw data for both methods, and demonstrate an overall superior performance of the ITRAP method in terms of both data consistency and performance. In conclusion, this study demonstrates that Improving the data quality by optimizing signal yield from high throughput studies of TCRpMHC-specificity is paramount in increasing our understanding of T cell-mediated immunity.

## Introduction

The specificity of T cells form the hallmark of cellular immunity. T cell specificity is determined by the triad of interactions between the T cell receptor (TCR), a peptide (p), and its restricting major histocompatibility complex (MHC). The TCR is a heterodimeric protein, typically consisting of an α- and β-chain. These chains are formed during T cell development as a result of stochastic recombination of the Variable (V), Diversity (D) and Joining (J) genes (1–5). As a result of the somatic recombination, highly variable joining segments are introduced, VJ and VDJ for α- and β-chains respectively, facilitating a diverse TCR repertoire that ensures protection from a broad and ever-changing range of pathogens or cancerous mutations (6,7). The joining segments are contained in a region known as the complementarity determining region 3 (CDR3). CDR1 and CDR2 reside in highly polymorphic regions of the V-gene. These three CDRs per chain form flexible loops of the TCR which engage with the peptide-MHC (pMHC) complex, thereby determining the specificity of the T cell (1–5,8,9).

Recent studies have elucidated shared TCR sequence features of T cells that share a common specificity, and for selected pMHCs, it has been possible to train models allowing to predict binding for TCRs novel to the trained model (1–5,10,11). The current primary limitation for the further development of such models is the lack of training data in terms of both quantity and diversity. Traditionally, TCR specificity data have been generated by assays such as multimer sorting and re-exposure assays, followed by bulk sequencing of typically the CDR3-loop of the TCRβ-chain. Such approaches hence fail to provide information about the paired TCR α- and β-chains. Recent studies have demonstrated that such paired information is essential to properly deduce and model TCR specificities (12,13). However, the advent of single-cell sequencing platforms promises a solution to this, generating high-throughput paired α-/β-chain TCR data. In addition these platforms intrinsically provide information on both positive as well as negative binding pairs (14), which is crucial when training machine learning models.

10x Genomics has specifically developed an immune profiling platform that couples TCR sequencing of both α- and β-chains with DNA barcoded peptide-MHC (pMHC) multimers, DNA barcoded surface marker antibodies, and DNA barcoded cell hashing antibodies. The platform is designed to capture a single cell together in a gel-bead in emulsion (GEM) (15,16). Each GEM contains GEM-specific barcoded primers which ensure the back-tracing of transcripts to the cell of origin. As the platform promises single-cell capture, the contents of a GEM should reflect a single cell and its associated barcoded analytes, hence GEM and T cell may be used interchangeably. The GEM primers also contain a unique molecular identifier (UMI) which ensures the quantification of transcripts unbiased by PCR amplification(17). Thus, single-cell screening of TCR-pMHC interactions yields the sequences of the TCR α-/β-chains and the expression level of both chains as well as the count of each unique pMHC binding event which can be interpreted as a proxy for T cell binding affinity (14).

In 2019, 10x Genomics released a large, state-of-the-art data set (14) which spurred activity within the TCR-pMHC modeling community (1–5,14,19, 25). The 10x Genomics data contained T cell specificities from four healthy donors screened against a panel of 50 pMHCs which includes 44 pMHCs for positive selection and 6 negative control pMHCs (14). However, this data presented new challenges. The single-cell platform is generally associated with a poor signal-to-noise ratio due to GEM-to-GEM leakage and capture of ambient analytes from suspension. The challenge was handled in various ways. In NetTCR-2.0, the data was utilized solely to define negative TCR-pMHC pairs, i.e. pairs that were not detected to bind any of the investigated pMHC complexes, thereby bypassing handling the noise within the detected positive data (12). Since the true TCR-pMHC pairs are a point of contention, the authors of ImRex purposefully omitted the 10x data (18), while the authors of TcellMatch and DeepTCR relied on the network to extract the salient pMHC-specific features of the TCRs (19,20). The authors of TCRAI were the first to develop a computational method, named ICON (Integrative COntext-specific Normalization), to discriminate true TCR-pMHC binding signal from nonspecific background noise (21). ICON was developed based on 10x Genomics data, utilizing the negative controls to empirically estimate the background binding noise per donor. The UMI counts of pMHCs were then corrected by subtracting the donor-specific estimated background noise. UMI counts were further corrected by penalizing pMHCs multiplets i.e., GEMs containing multiple DNA barcodes corresponding to two or more different pMHCs. The final step of ICON is the normalization of UMI counts across pMHCs and GEMs to make them directly comparable. As a result, ICON identified a total of 53,062 T cells belonging to 5,722 unique clonotypes.

Recently, we have proposed an alternative denoising framework: ITRAP (improved T cell and Antigen Pairing) (22). The ITRAP framework was originally developed and tested on in-house single-cell data generated using the 10x Genomics platform similar to the public 10x Genomics data. In comparison to ICON, ITRAP takes a different approach for denoising. The key is to study GEMs in an ensemble rather than individually, since this allows deviations to be averaged out. That is, If a pMHC is distributed with a significantly higher mean UMI in the ensemble compared to others, we expect this pMHC to reflect the true target of the clonotype, collectively providing a golden standard. The ITRAP framework consists of a series of filtering approaches to obtain increasingly accurate TCR-pMHC pairing. The first filtering step is based on identifying expected targets by comparing the UMI distributions of all pMHCs detected within a clonotype consisting of 10 or more GEMs. Based on the labeling of true and false targets, an accuracy score is next defined, and thresholds on UMI counts can be defined to maximize this accuracy. By globally applying the optimal threshold, the remaining clonotypes are next filtered to ideally represent the same level of accuracy in their pMHC annotations. Another key step of ITRAP filtering is ensuring HLA correspondence between pMHC and the HLA haplotype of the T cell donor. In immune profiling assays, the option to hash cells by donor-of-origin enables the assignment of HLA haplotype restriction to each cell. Correspondence between the allele of pMHC and donor haplotype can be used to verify the assignment of the pMHC, assuming that a T cell is restricted solely to the allele for which it was selected during the thymocyte maturation process. In the public 10x data, the cells are not hashed, however, the experiment was run in parallel for each donor, enabling *in silico* hashing of the individual single-cell runs.

In this study, we report a benchmark of the ICON and ITRAP frameworks. Both methods are applied to the 10x Genomics data since this is the only data set containing negative controls as is required by ICON. As no external golden standard exists, the performance of the two methods is evaluated on internal performance metrics GEM retention, accuracy, average binding concordance, and AUC of similarity scores earlier presented by Povlsen et al. (22), as well as in terms of predictive performance of machine learning methods trained on the raw and denoised 10X data on independent data.

## Material and Methods

### Data Retrieval

The 10x Genomics data set used for this study was downloaded from https://support.10xgenomics.com/single-cell-vdj/datasets (Application Note. A New Way of Exploring Immunity).

The ICON-filtered dataset was curated by Zhang et al. employing ICON for identifying reliable TCR-pMHC interactions. The resulting filtered data was downloaded from http://advances.sciencemag.org/cgi/content/full/7/20/eabf5835/DC1. This dataset contains 53,062 cells (here referred to as GEMs) that passed the ICON filtering with ICON-corrected pMHC and TCR annotations. The ICON output provided with the publication contains a fifth donor, donor V, which was removed from the set (14,052 GEMs). This donor V was part of an internal experiment by Zhang et al., and raw data was not available and has not been acquired for this benchmark.

### Data Curation

The data consists of four sets of single-cell RNA sequencing and immune profiling from four healthy donors. The sets were concatenated for one combined analysis. GEM-specific 10x barcodes (GEM barcodes) were observed in duplicates across the donor sets (625,170 duplicate GEMs of raw TCR annotation and 386 duplicate GEMs of raw pMHC annotations). As a result the GEM barcodes were additionally suffixed by donor, i.e.AAACCTGTCTAACTTC-6-s2. Cells (referred to as GEMs) were removed if the annotated CDR3αβ sequences were not productive, full length, or contained non-IUPAC characters, resulting in 181,913 GEMs.

### ITRAP Data Filtering

ITRAP consists of different types of filters that can be applied to single-cell immune profiling data to reliably identify TCR-pMHC interactions. The accepted inputs include single-cell RNA sequencing, targeted T cell receptor sequencing, dCODE-Dextramer sequencing for DNA barcoded pMHC multimers, as well as CITE-seq sequencing of DNA barcoded cell hashing antibodies. The method includes the following major steps as described in (22) and applied to the current data as outlined below (note that the filtering process is subsequent and can be stopped at any given user-defined step):

Step 1: *Correction of 10x annotated clonotypes.* Instead of limiting clonotypes to groups of GEMs with exact nucleotide sequence identity, clonotypes were defined based on VJαβ-gene annotation and the CDR3αβ amino acid sequences. For clonotypes for GEMs containing only one TCR chain, the other chain was imputed if the present chain matched only one pre-established clonotype. GEMs containing multiple chains were annotated by the most abundant chain by UMI count.
Step 2: *Filtering based on data-driven thresholds.* For clonotypes consisting of more than 10 GEMs, the expected target was identified if a pMHC had a significantly higher UMI distribution than other pMHCs also captured in GEMs of the given clonotype. Significance was tested by Wilcoxon at α=0.05. The pMHCs not declared as target are considered background noise. An accuracy score was obtained based on the fraction of target pMHCs over background pMHCs. The optimal UMI threshold was selected as the UMI value that maximized this accuracy score. Next, the thresholds were applied to the entire data set, and for each retrained GEM, the pMHC target was assigned from the highest pMHC UMI count.
Step 3: *Match pMHC HLA allele with donor haplotype.* The HLA-A, -B, and -C haplotypes were provided by an application note following the release of the single-cell sequencing of the four healthy individuals. Since the samples were sequenced individually the haplotypes were easily added to the data sets. GEMs consisting of a mismatch between donor haplotype and pMHC were discarded.
Step 4: *Selecting GEMs with paired αβ chains.* GEMs with only a single chain were removed. For GEMs with multiple α- and/or β-chains, the ones with the highest UMI counts were assigned to each GEM.
Step 5: *Filtering specificity singlets.* If a TCR-pMHC pair was only observed once, it was discarded to increase confidence in matches.
Step 6: *Selecting 10x annotated cells.* Application of the 10x provided filter “is_cell” (14).

### TCR Specificity Prediction

In order to quantify the benefit of removing noisy observations from the original 10x dataset, we trained the NetTCR-2.1 CDR3αβ framework on i) the unfiltered 10x data, ii) the ITRAP-filtered data using optimal UMI threshold and donor HLA matching, iii) ICON-filtered data, with the setup recommended by the authors (21). For details on the modeling framework refer to (23). In short, NetTCR-2.1 uses convolutional neural networks to predict the binding of a TCR and a peptide-MHC complex. In the current work, the inputs to the model are the CDR3 α and β amino acid sequences. For each of the peptides present in the data set, a model is trained on the TCR data specific to that epitope. These models are subsequently used to obtain predictions over an external evaluation set.

Both the raw and filtered data consist of GEMs with annotated TCRpMHC triads. In each case, a set of positive TCR-peptide pairs was built by selecting, for each clonotype, the most frequent pMHC across GEMs (for that clonotype) as the target pMHC. To validate the trained models, an external evaluation set was retrieved from VDJdb (24). This dataset consisted of 927 TCR sequences relative to 4 epitopes (GILGFVFTL, GLCTLVAML, ELAGIGILTV, IVTDFSVIK). Also, the training set was restricted to the set of 4 peptides, to ensure overlap between the training and evaluation set. For both data sets, negative peptide-TCR pairs were artificially generated by pairing the positive TCRs with the other 3 peptides different from their target cognate.

To investigate performance inflation due to a similarity overlap between training and evaluation sets, TCRs from the evaluation data that had a kernel similarity value (25) above 0.9 to the training TCRs were removed.

The training set was randomly split into 5 partitions and the models were trained using 5-fold nested cross-validation. The resulting 20 trained models were used in an ensemble to get predictions over the TCRs in the evaluation set. The different training and evaluation datasets are available at https://github.com/mnielLab/ITRAP_benchmark.

## Results

### Summary of the public 10x data

The public data set made available by 10x Genomics is the result of screening CD8^+^ T cells from four healthy donors against a panel of 50 pMHC DNA barcode-labeled multimers. The complete data was initially reduced to only include IUPAC encoded amino acids within CDR3 sequences and further only considered GEMs which contained both TCR and pMHC annotations, resulting in 181,913 GEMs. Donors investigated by 10x Genomics were selected by HLA haplotype to ensure overlap with the HLA alleles of the pMHC panel. 44 of the multimers contain antigenic peptides derived from CMV, EBV, influenza, HTLV, HPV, HIV, and known cancer antigens. It should be noted that the donors were all seronegative for HIV, HBV, and HBC. The remaining six multimers contained negative control peptides restricted by five HLAs. The specificities of each of the four donors were screened in parallel i.e., in four different experimental runs. Therefore, unique GEM-specific 10x barcodes (GEM barcodes) were in some cases observed in replicas across runs. In order to distinguish these distinct GEMs, an extra suffix was added denoting the donor (sample ID). The unfiltered output is portrayed in Figure 1, which clearly demonstrates the issue of noise, as every GEM contains multiple pMHCs. Most GEMs contain TCRs annotated with a unique α- and β-chain, however, ~16% are annotated with only an α- or a β-chain while ~10% are annotated with multiple α- or β-chains, which further challenges the investigation of specificity.

**Figure 1:**
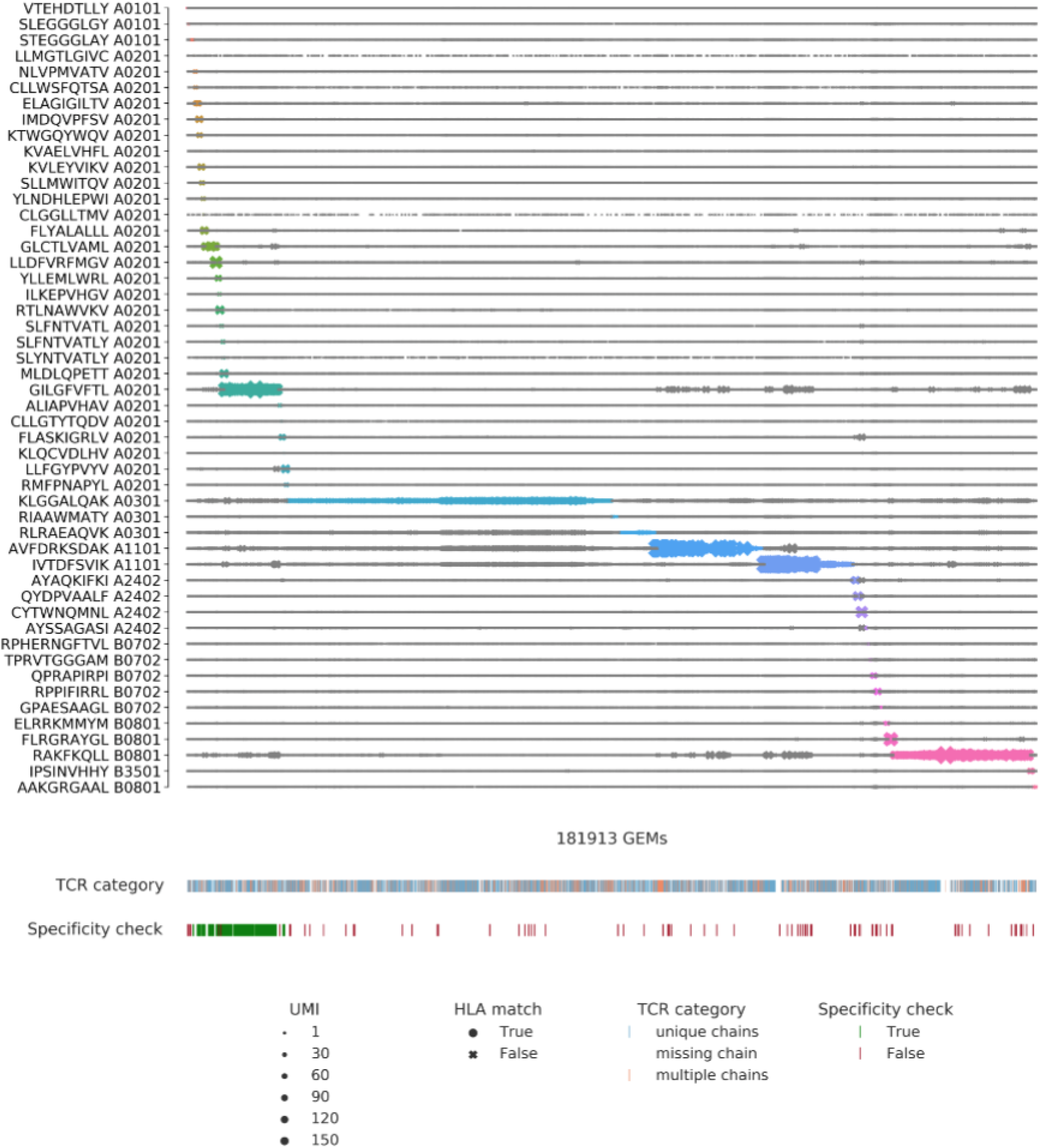
Visualization of all detected pMHC barcodes (y-axis) within each of the 181,913 GEMs (x-axis). In each GEM the most abundant pMHC is marked by a color, while the remaining pMHCs in the GEM are gray. The marker size reports the UMI count of the given pMHC and the shape recounts whether the HLA allele of the pMHC matches the HLA haplotype of the donor, which is provided in the experimental report (14). The first color bar indicates the type of TCR chain annotation; whether the TCR has a unique αβ-pair, is missing a chain, or consists of multiple chains. The second color bar is a specificity check against the specificity databases IEDB and VDJdb. Colors highlight the GEMs where the CDR3αβ sequences are contained in the databases. The green color represents a match between the database pMHC and the detected pMHC, while red indicates a mismatch.

### Alignment of ICON- and 10x-assigned GEMs revealing inconsistent annotations

In order to compare the ICON and ITRAP filtering frameworks, the outputs from each method were aligned based on the GEM barcode, consisting of 16 nucleotides, a suffix pertaining to the sequencing well, and a sample ID suffix. ICON reported retention of 53,062 GEMs out of the total set of 181,913 GEMs. However, ICON only contains 5,031 GEMs that match the original database on the full GEM barcode, due to inconsistencies in the suffix annotation. When stripping the barcode down to only the 16 nucleotides, we were able to align 39,806 GEM barcodes, as exemplified in Figure 2a. We also observed inconsistencies of TCRαβ annotations in 3,391 GEMs, as illustrated in Figure 2b+c. 1,854 GEMs were missing either an α- or a β-chain in the 10x data, but not in the ICON set, while 1,537 GEMs were fully annotated, but had inconsistent TCR annotations between ICON and the 10x data. The inconsistencies in TCRαβ annotations may have arisen from imputations based on the 10x-provided clonotype summary. However, such imputation should be performed with caution because the same CDR3 may form part of several different clonotypes. The example given in Figure 2b represents an imputation likely based on the CDR3β sequence. In this example, the CDR3β sequence is part of 42 distinct 10x clonotypes, all carrying the same CDR3β sequence, but paired with different CDR3α sequences. The same case is made for 2c and all the other inconsistent GEMs.

**Figure 2:**
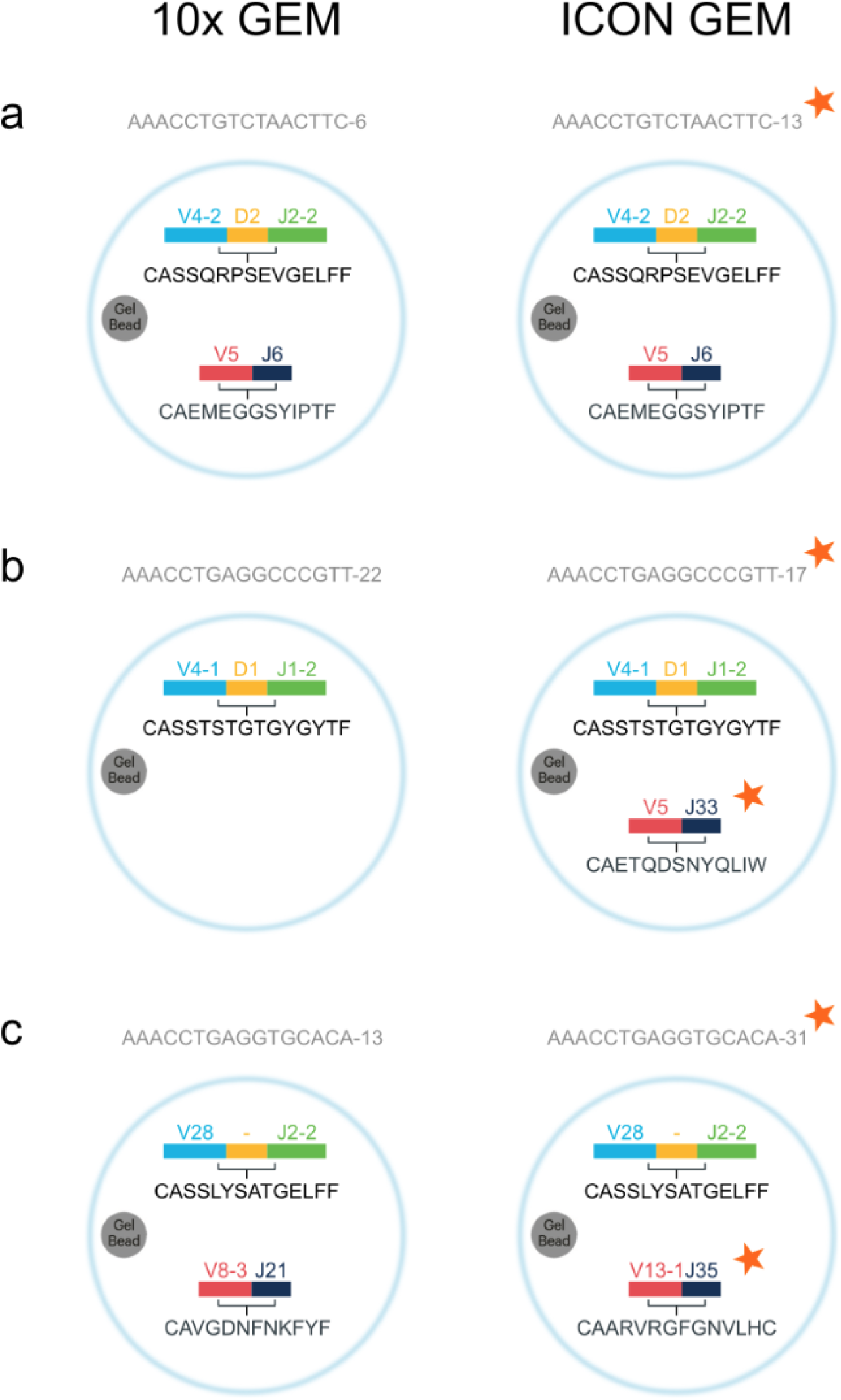
Illustrations of annotation inconsistencies. The figure shows examples of GEMs and their TCR annotations from 10x and ICON, respectively. The observed inconsistencies are grouped into three major groups. The inconsistencies are highlighted with a red star in each group. (a) 33,342 GEMs were mapped from the ICON set with inconsistent GEM barcode suffixes. Mapping was based on the GEM barcode nucleotide sequence and TCR annotations. (b) 1854 GEMs were missing either an α- or a β-chain in the 10x data, but not in the ICON set. (c) 1537 GEMs were fully annotated, but the TCR annotations were inconsistent between ICON and the 10x data.

Imputation by 10x clonotypes is further made difficult as their clonotype definition allows multiple α- or β-chains in one clonotype, perhaps a reflection of incomplete allelic exclusion. Thus, 116 of the fully annotated GEMs with mismatching TCRαβ annotations between ICON and 10x can be explained by a switch from one chain to the other, still within the same clonotype definition. This non-conformity has challenged the benchmark, however, we have proceeded assuming that there is a reasonable, however undocumented, explanation for GEM assignments provided in the ICON data set.

### ITRAP - Revisiting clonotype assignment

For efficient utilization of ITRAP, the 10x-assigned clonotypes were redefined. The original annotations of clonotypes were based on unique nucleotide sequences of the T cell receptor to identify expansions of clonally-related T cells. However, the somatic pedigree is not relevant for understanding the biochemical properties of the TCR. Instead, we are interested in grouping T cells of TCRs with identical amino acid sequences including identical CDR3s. This regrouping of GEMs results in larger clonotypes beneficial for statistical power in the ITRAP filtering steps. Thus, in ITRAP, a clonotype is generally defined by a unique set of Vαβ- and Jαβ-genes as well as CDR3αβ, with few exceptions of repeated chain multiplets across GEMs. It should be noted that redefining clonotypes does not affect the individual GEM annotations of Vαβ Jαβ-genes or CDR3αβ sequences but only pertains to how GEMs are grouped and labeled. Redefining 10x clonotypes resulted in 76,627 unique clonotypes.

### The optimized ITRAP threshold on UMI counts

Of the 76,627 clonotypes, 1,151 were represented by 10 or more GEMs, and for 1,107 of them, we were able to annotate an expected binder (for details on this step refer to Materials and Methods). Running the UMI threshold search step on these clonotypes optimizing the proportion of GEMs where the most abundant pMHC aligns with the expected target, the derived optimized thresholds values were: a minimum UMI of 5 for any pMHC. For pMHC multiplets, the most abundant pMHC must be 1.2 times greater in UMI counts than the second most abundant pMHC, and a minimum of 1 UMI for TCR α- and β-chains. By this filter, the data set is reduced to 91,652 GEMs and 27,925 unique clonotypes. Additionally, filtering on matching HLA serves as the recommended minimum of filters for ITRAP. Doing this results in a set of 40,584 GEMs and 6,751 unique clonotypes with complete TCR annotation. These stepwise filtered data are available from https://services.healthtech.dtu.dk/suppl/immunology/ITRAP_benchmark/.

### Benchmark of ICON and ITRAP

The two filtering frameworks were benchmarked on four performance metrics, as described by Povlsen et al. (22): fraction of retained GEMs, accuracy of specificity, average binding concordance across all clonotypes, and AUC of CDR3αβ similarities. Accuracy is computed as the fraction of GEMs where the most abundant pMHC (by UMI counts) corresponds to the expected binder of a clonotype. An expected binder is defined for each clonotype as the pMHC which is distributed with a mean UMI count significantly higher than all other pMHCs detected as binders for the given clonotype (Wilcoxon, α=0.05). Binding concordance is computed as the fraction of GEMs within a clonotype that binds a given pMHC and describes the dispersion of pMHC annotations within the clonotype. In a data set where no cross-reactivity is expected, excluding technical artifacts, the average binding concordance should be 100%. Finally, the similarity between two TCRs is defined as the summed score of the pairwise CDR3α and CDR3β similarities each calculated using the kernel similarity method described (25). Here, in short, clonotypes within a given “plateau” (i.e clonotypes annotated to a given peptide) were compared against other clonotypes within the plateau and the same number of randomly sampled clonotypes from other plateaus, and a maximal inter and intra similarity score found. The AUC metric is computed based on the hypothesis that different TCRs binding the same pMHC (intra-specificity) are more similar to each other than to TCRs of other specificities (inter-specificity) (26).

The summary of both filtering frameworks across our selected performance metrics is presented in Figure 3. Both ICON and ITRAP discard a large number of GEMs (Figure 3a). The recommended filtering steps for ITRAP consist of filtering on UMI thresholds and matching HLA between annotated pMHC and HLA haplotype of the donor, which yields 40,584 GEMs, which is slightly more than ICON (39,806). This is, both methods discard ~78% of the raw input data. In terms of performance metrics, the results reveal an overall high performance of both frameworks.

**Figure 3:**
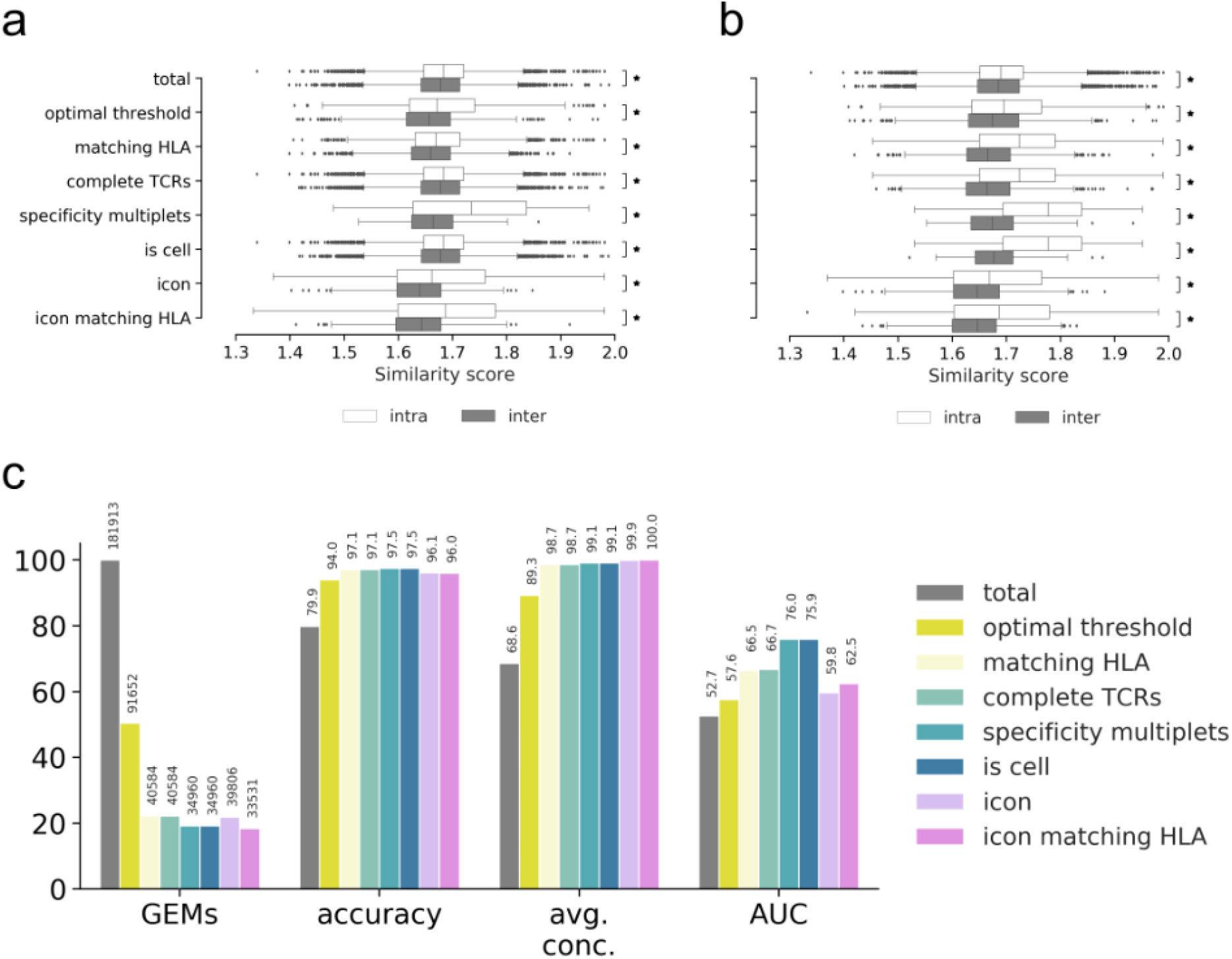
Performance metrics for evaluating the filtering steps of ITRAP with ICON. The ITRAP filtering steps consist of total (raw, unfiltered data), optimal threshold obtained from grid search, matching HLA, complete TCRs with a unique set of α- and β-chain, specificity multiplets i.e., TCR-pMHC pairs observed in two or more GEMs, and “is cell” defined by 10x Genomics Cellranger. ICON yields a single output, however, an addendum has been made to also filter ICON output on HLA match between pMHC and HLA haplotype of the donor. (a) The boxplots show kernel similarity scores between CDR3β sequences of intra- (white) and inter- (dark) specificity for each of the filtering steps. A significant difference (Wilcoxon, α=0.05) of mean between inter- and intra-specificity is marked with an asterisk to the right (b) (for details on this metric refer to text). Here, the boxplots show the cumulative effect of ITRAP filters on similarity scores. (c) Performance is measured and summarized by a number of metrics: ratio of retained GEMs (GEMs), accuracy defined by the proportion of GEMs where most abundant pMHC matches the expected binder (accuracy), average binding concordance (avg. conc.) and AUC of similarity scores (AUC). The ITRAP filters are also here cumulatively added to show increasing improvement in performance.

In terms of accuracy, both methods achieve a performance gain of ~22% (improved from ~80% to 97%) with a slight advantage in favor of ITRAP. When it comes to concordance, a similar large gain was observed for both methods with an advantage in favor of ICON. This is a consequence of the fact that ICON was essentially designed for optimizing this metric. Figure 3b and 3c show the effects of each filter either alone (b) or in combination (c), in terms of the distribution of intra-versus inter-specificity distributions. The results clearly demonstrate how the separation of inter- and inter-specificity improves as more filters are applied. To quantify the separation of distributions, an AUC score was computed from the principles that perfect intra-specificity scores are close to a maximum value of 2, while inter-specificity resembles completely different TCRs of similarity close to 0. Note that AUC here does not translate into a predictive performance, but rather reflects the extent to which intra-similarity can be distinguished from inter-similarity values. These AUC values are displayed as the last performance metric in Figure 3a, again confirming an improved quality of the data after filtering by both methods. Applying an additional specificity multiplet filtering step removing specificity singlets, removes an additional small set of 5624 GEMs, but yields a high gain in AUC and results in the largest separation between intra- and inter-specificity distributions of all filtering steps. Here, a specificity singlet is defined as a TCR-pMHC pair only detected with a single GEM, which makes the pairing more susceptible to artifacts. However, those GEMs represent unique clonotypes, so this filter also vastly reduces the total number of clonotypes.

As mentioned, ICON does not discard GEMs based on HLA match between pMHC and donor haplotype. However, we have tested the impact of adding that filter to ICON, which reduces the yield to 33,531 GEMs. Based on the AUC of similarity scores, the recommended ITRAP filters yield a slightly higher performance compared to ICON (66.5 versus 62.5).

In conclusion, ITRAP overall outperforms ICON in these benchmark evaluations both in terms of yield, accuracy and similarity AUC.

### Visual inspection of ICON and ITRAP outputs

The differences in binding concordance between ITRAP and ICON are clearly visualized in Figure 4 and Figure 5. Figure 4 presents the ITRAP-filters of UMI threshold, HLA matching, and complete TCRs i.e., unique pairing of α- and β-chain.

**Figure 4:**
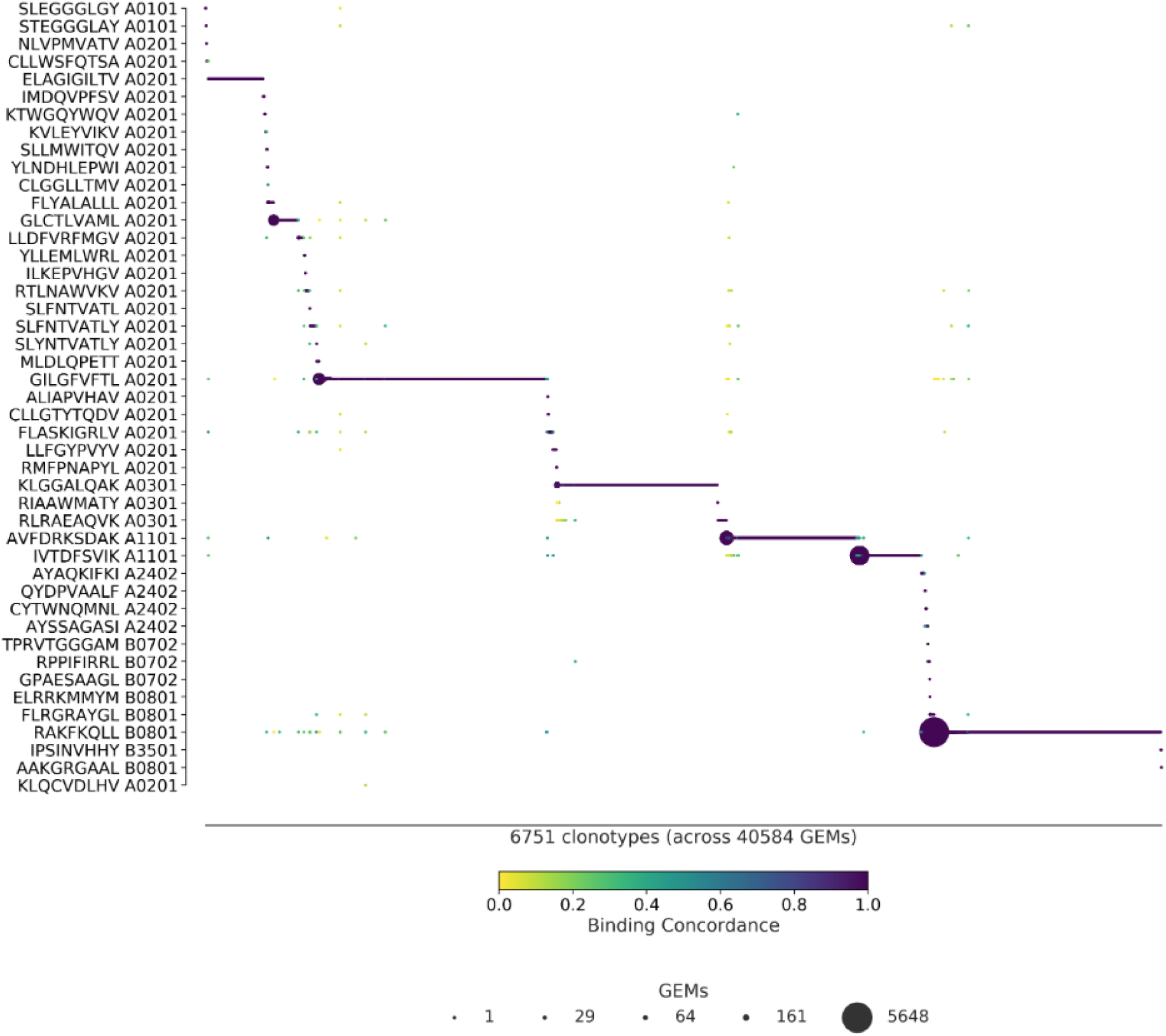
ITRAP-derived specificity per clonotype. ITRAP-filters consist of UMI threshold, HLA matching, and complete TCRs i.e., a unique pairing of α- and β-chain. The library peptides are listed on the y-axis and each clonotype is represented on the x-axis. Below the x-axis is annotated the total number of clonotypes and GEMs in the presented data. The marker size shows the number of GEMs supporting a given specificity. The color indicates the binding concordance which is calculated as the fraction of GEMs within a clonotype that supports a given pMHC. The higher the concordance, the larger the fraction of supporting GEMs.

**Figure 5:**
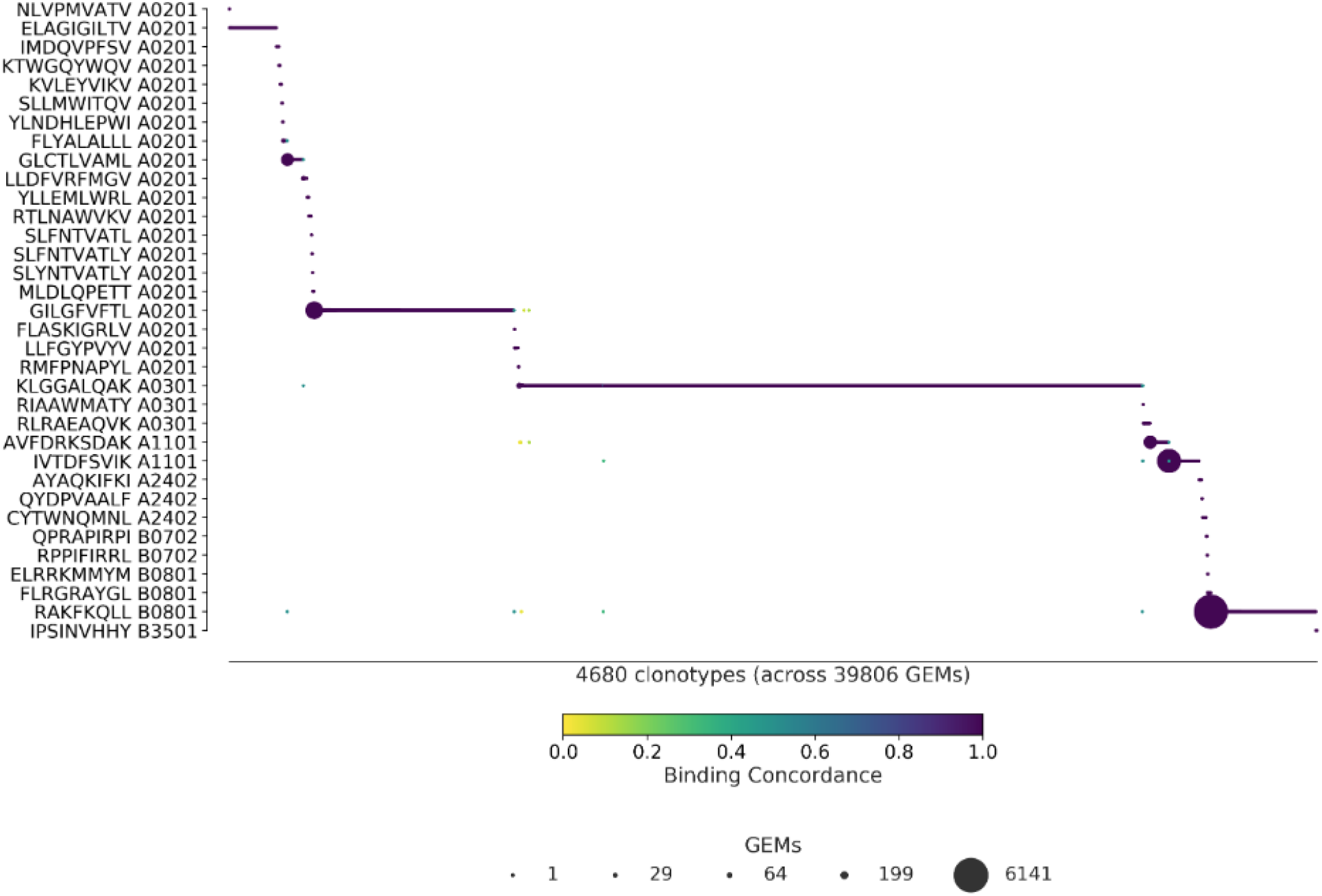
ICON-derived specificity per clonotype. The library peptides are listed on the y-axis and each clonotype is represented on the x-axis. Below the x-axis is annotated the total number of clonotypes and GEMs in the presented data. The marker size shows the number of GEMs supporting a given specificity. The color indicates the binding concordance which is calculated as the fraction of GEMs within a clonotype that supports a given pMHC. The higher the concordance, the larger the fraction of supporting GEMs.

With an average binding concordance of 98.7, we observe 407 GEMs with a binding concordance of less than 50%, which we will refer to as outliers. A substantial proportion of these are pMHC targets annotated to different HLA alleles. This contradicts the prevailing belief that T cells are restricted to the HLA for which they were positively selected during maturation. We thus suspect that some of these events are a result of random capture of ambient multimer barcode.

In 65 GEMs of the 407 outliers, an expected pMHC target had not been identified, due to the small sizes of the clones. Of the remaining 320 outliers, 76 GEMs exhibit a pattern that aligns with potential cross-reactivity. That is, a TCR will typically have a single, preferred target while allowing binding of other pMHCs to a lesser extent, i.e. clones of a clonotype may display a single dominant pMHC response of high binding concordance with few smaller responses of low binding concordance. For the clonotypes of these 76 GEMs, the dominant high-concordance pMHC coincides with the expected target of the individual clonotypes. In 18 of these GEMs, the corresponding clonotypes showed divergent HLA restriction between the annotated low-concordance pMHC and the expected target for the given clonotype. In all of the 76 GEMs, the expected target was detected albeit at a lower UMI count than the annotated pMHC.

The remaining set of 266 GEMs consists of 80 clonotypes exhibiting highly dispersed binding to many different pMHCs, all with low binding concordance. All of these GEMs also contain multiplets of pMHCs. Based on these observations, we conclude that the majority of the 407 outliers are likely artifacts that have escaped the ITRAP filtering steps and thus not true cross-binding events.

Figure 5 presents the ICON retrieved specificities. With an average binding concordance of 99.9%, most clonotypes are paired with a single specificity, and only 24 GEMs are categorized as outliers. 13 of the outliers are annotated with a pMHC that does not match the allele of the donor. 4 of the outliers contain CDR3 sequences that differ from the 10x annotation and may be a result of imputation.

Finally, a key difference between the two methods is that ITRAP retains 45 pMHCs from the staining whereas ICON retains 34 pMHCs. The 11 peptides retained by ITRAP and not ICON elicit small and few responses, but are primarily not involved in cross-binding events. With both filtering frameworks, the largest responses are toward KLG HLA*A-03:01 (n=1085 for ITRAP, n=2681 for ICON), RKA HLA*B-08:01 (n=1178 for ITRAP, n=450 for ICON) and GIL HLA*A-02:01 (n=1337 for ITRAP, n=842 for ICON). ICON retains more GEMs and more clonotypes within these peptides, at the expense of other specificities, than ITRAP does.

### Predicting TCR specificity with ITRAP- and ICON-filtered data

To quantify the potential predictive performance gain derived from filtering and denoising the raw TCR data, we trained NetTCR-2.1 (12) on the raw 10x data and on the ICON and ITRAP-filtered datasets (for details refer to Materials and Method). Note, that the data split for training here was done randomly for the three data sets, likely inflating the reported cross-validation performance. However, since the aim is to compare the methods and not create a new model, we deem this justified. We evaluated the performance of the three models on an independent dataset derived from VDJdb (24). The evaluation set consisted of 927 positive TCRs relative to the 4 peptides in consideration. Table 1 reports the number of positive evaluation TCRs for each epitope. The counts of positive TCRs are reported both before and after similarity filtering. Negative data was added by introducing swapped TCRs as described in materials and methods.

**Table 1:**
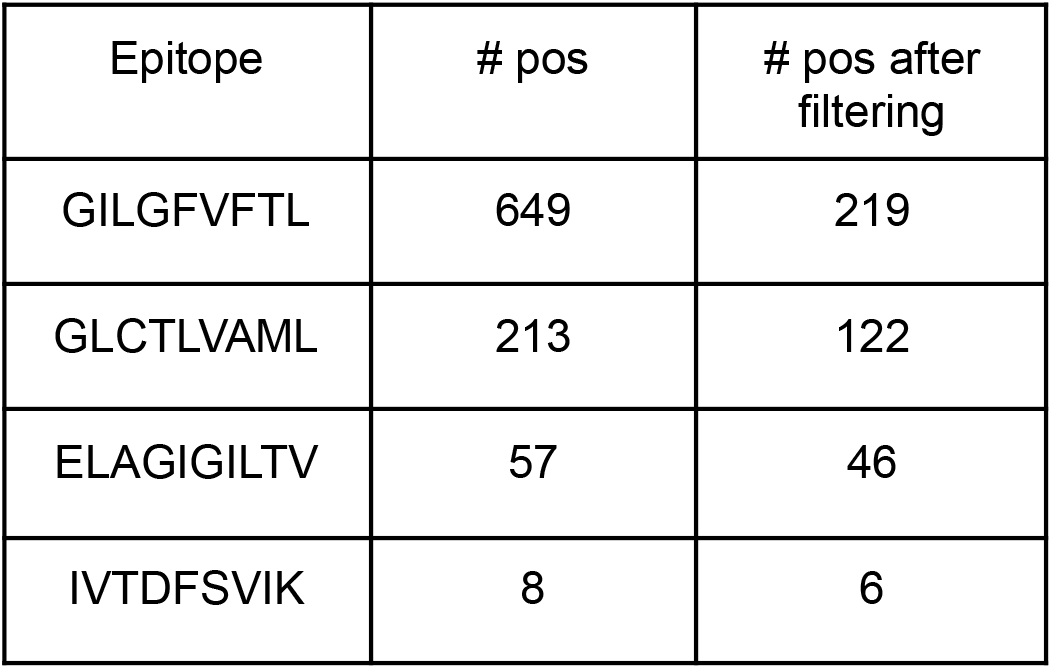
Counts of positive TCRs in the evaluation set, for each peptide. The first column contains the epitope sequence, the middle column contains the count of positive TCRs in the original evaluation set, the last column reports the number of positive TCRs left after the similarity reduction step.

| Epitope | # pos | # pos after filtering |
| --- | --- | --- |
| GILGFVFTL | 649 | 219 |
| GLCTLVAML | 213 | 122 |
| ELAGIGILTV | 57 | 46 |
| IVTDFSVIK | 8 | 6 |

The results of the experiment are shown in Figure 6. The cross-validation performance refers to the performance on the concatenated test sets while the predictions on the evaluation set were calculated as an ensemble of the predictions of the 20 trained models. For the evaluation predictions, we reported the AUCs on the full evaluation set (middle panel) and on the similarity reduced set (lower panel). For each trained model, the AUC was reported on a per-peptide level. An overall performance value was also given by averaging AUCs across peptides. We reported the average AUCs both as a mean value of the AUCs from each peptide and as a weighted average of the peptide AUCs, weighted by the number of positive TCRs for that specific peptide in the dataset.

**Figure 6:**
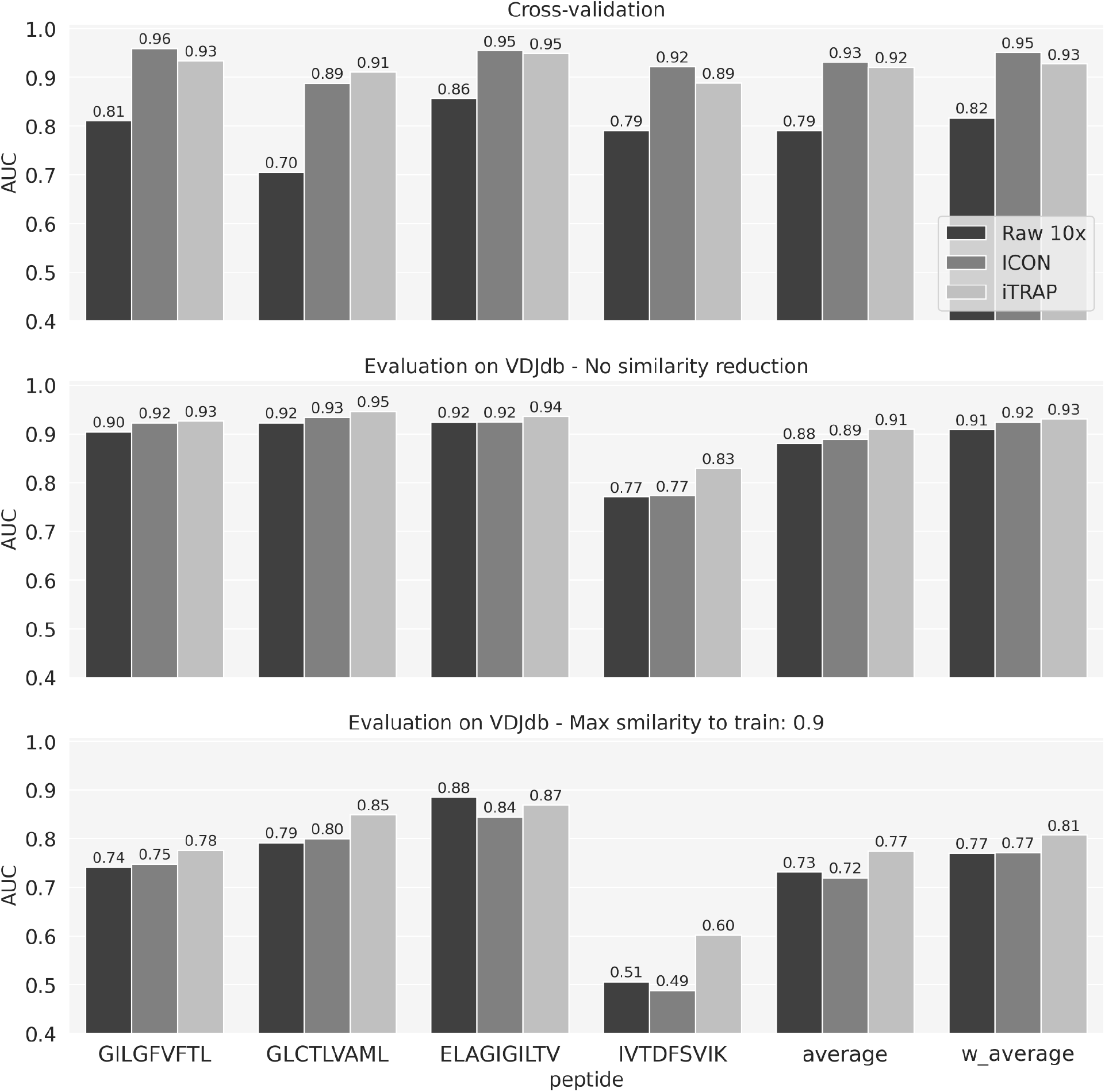
Performance of NetTCR-2.1 in terms of AUC on the raw 10x data and on the filtered datasets. The AUC is given on the concatenated test sets from cross-validation and on the external evaluation set from VDJdb (before and after removing evaluation TCRs similar to sequences in the training set). ‘average’ refers to the mean of the AUC values across peptides; ‘w_average’ is a weighted average of AUCs across peptides, weighted by the number of positive TCRs for the peptides in the dataset in consideration.

Both in cross-validation and on the external data, the models trained on ICON and ITRAP datasets outperformed the models trained on unfiltered data. Interestingly, the ICON outperformed ITRAP on almost all the peptides in cross-validation. This can be explained by looking at the similarity between the test and training partitions in cross-validation. Figure 7 shows, for each peptide, the distribution of kernel similarities between the positive TCRs in the test set and their nearest neighboring positive TCR in the training. For the GIL and IVT peptides, ICON has a higher median similarity between training and test set, leading to a higher AUC value in cross-validation for these two peptides. The models trained on ITRAP-filtered data generalize better on the external dataset, outperforming ICON across all peptides. For the similarity-reduced evaluation set, all the differences in AUC between ICON and ITRAP are significantly different for all peptides except IVT (p<0.05, bootstrap test on the AUC with 100 repetitions). This is also confirmed by the improved average and weighted average performance of the model trained on ITRAP data. Furthermore, the gap in performance between the two methods was increased when the overlap between the training and evaluation set was reduced. ITRAP showed a 2% increase in average AUC (and 1% weighted average AUC) compared to ICON on the evaluation set with no similarity filtering; when removing the evaluation TCRs with a similarity >0.9 to the training set, ITRAP yielded 5% gain in average AUC (4% weighted average AUC).

**Figure 7:**
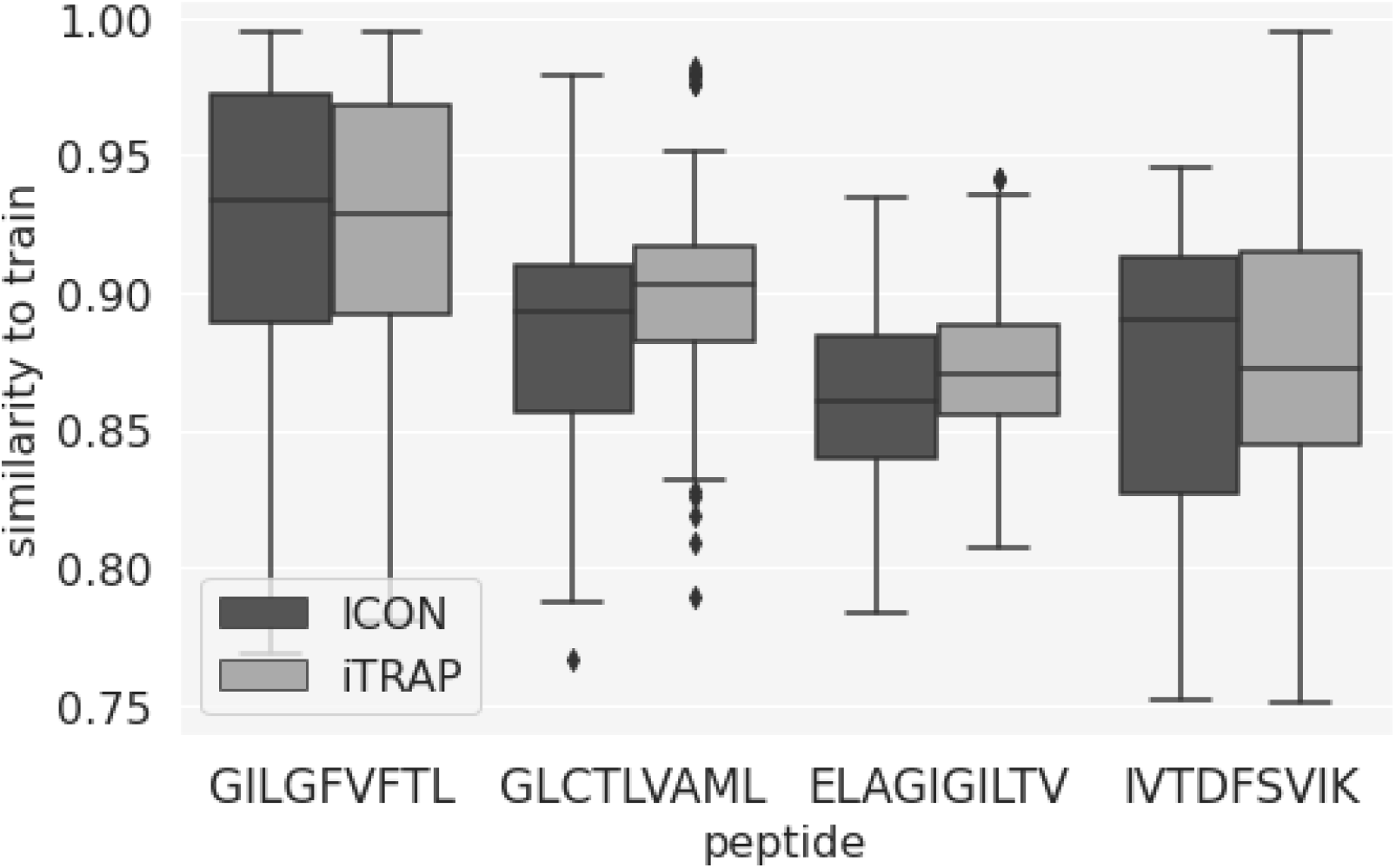
Kernel similarity values between positive TCRs in the test set and their nearest neighbor in the positive set of training TCRs.

## Discussion

T cell specificity is defined from subtle sequence and structural properties of the TCRs, and learning the rules defining this specificity has been challenged by the current lack of a large number of observations of high-quality paired TCRαβ data with known pMHC targets. Single-cell screening assays may pave the way to resolve this. The technology enables the study of TCR-to-pMHC binders, decisive non-binders, and even cross-binding. However, single-cell data is generally characterized by low quality, and denoising single-cell specificity data is therefore a critical bottleneck in studying T cell specificity. Here, we have evaluated two methods, ITRAP and ICON, both aiming at resolving this bottleneck, filtering noise and putative artifacts from true binding events. Since no golden standard exists, the methods are evaluated via metrics designed for the purpose, as well as in the application of developing prediction models for TCR specificity, enabling data-driven quality assessment.

Both methods were applied to the same raw dataset generated by 10x Genomics. The two filtering frameworks both demonstrate very good performance, but with diverging upsides and downsides. ICON excels at reducing ambiguous specificity annotations, such that the majority of clonotypes are annotated with exactly one pMHC target. This however, also becomes a hindrance in detecting cross-reactivity. Conversely, the ITRAP method includes more GEMs across more pMHCs and does not explicitly exclude potential cross-reactive binding events.

The filtering frameworks were evaluated on four metrics: retention of GEMs, binding accuracy guided by expected targets, average binding concordance, and AUC of kernel similarity scores. Both methods discarded the vast majority, retaining only ~22% of the raw data. ITRAP achieved the highest accuracy score. However, binding accuracy may be a biased metric in this context as ITRAP was specifically designed to maximize this score. Similarly, ICON displayed a superior average binding concordance, favoring low dispersion of specificity within a clonotype, which ICON was purposefully designed to reduce. The AUC of kernel similarity scores used in this work as a metric for quantifying separation of distributions in terms of the TCR similarities within and between annotated clonotypes acts as the only method-independent metric. Here, ITRAP demonstrated improved performance compared to ICON.

Each framework has a set of requirements for the method to work optimally. ICON relies on gene expression data to remove duplicates and negative control pMHC multimers to correct binding signals of positive pMHC, and a set of negative control pMHCs to set a cutoff for pMHC UMI counts. The impact of gene expression data was previously tested for ITRAP, which showed only minute added performance (22). Due to the low impact and the high expense of running gene expression sequencing, this filtering step was deprioritized in ITRAP. ITRAP on the other hand heavily relies on cell hashing, where HLA typing of donors is known, to validate specificities (though the framework can be run excluding this filtering step). This, of course, also confers an additional cost. However, as demonstrated in this and the original ITRAP study, this step is critical when running the 10x analysis on samples for multiple donors, since it allows further denoising and analysis of donor specific T cell repertoires.

Both frameworks assume that the pMHC UMI count acts as a proxy for a given TCR-pMHC pair binding affinity, and use the count either directly (ITRAP) or corrected and normalized (ICON) to filter away GEMs. However, it is important to note that the UMI count refers to the number of pMHC multimers captured together with a T cell in a GEM. The count may be affected by the extent of ambient multimers, T cell expression of TCRs, and binding affinity/avidity. Thus to improve the filtering strategies of ITRAP or ICON future methods may implement adjusted TCR-pMHC pairing scores.

Pairing of TCR and pMHC is further made difficult in the cases where a presumed single cell expresses two different α- or β-chains. The dual expression is a known phenomenon (27)–(28), and thus cannot simply be written off as capture of multiple cells. Neither ICON nor ITRAP seek to investigate the impact on such specificities, but simply annotate the most abundantly expressed chain. To improve specificity detection, this aspect should be investigated further. Moreover, CDR3α- and β-pairs are not unique, but exist in various combinations, despite the stochastic process under which they are produced. Therefore, imputing CDR3 chains for GEMs with either multiple chains or GEMs missing a chain, will often not result in a unique pairing. We speculate that ICON has attempted such imputations since we observed discrepancies in CDR3 annotations between 10x and ICON data sets. The comparison was further complicated by inconsistent GEM barcodes between ICON and the 10x data. The alteration of barcodes is unaccounted for by the authors of ICON.

To further quantify how the two filtering approaches increased the signal-to-noise ratio, we trained TCR specificity prediction models on i) the raw 10x dataset, ii) the ITRAP-filtered data (using UMI threshold and HLA matching criteria), and iii) ICON-filtered data. The results showed that both ICON and ITRAP-filtered data sets lead to improved performance, compared to training on the raw 10x data. This further confirms that both methods filter out artifacts from the datasets, increasing the signal-to-noise ratio. The two models performed comparably in cross-validation. However, ITRAP demonstrated better generalizability compared to ICON on a novel set of TCRs obtained from VDJdb, independent from the training data.

## Conclusions

In conclusion, in this work we have compared two methods, ITRAP and ICON for denoising of single-cell TCR pMHC data. ITRAP was demonstrated to provide a higher yield (more GEMs and clonotypes covering more pMHC molecules were retrained), higher accuracy and large TCR similarity within and discrepancy between clonotypes compared to ICON. Using the data to train TCR specificity prediction models, also demonstrated a higher value of the ITRAP denoised data, resulting in models with higher generalization power compared to models trained on the ICON-filtered data. Moreover, and in contrast to ICON, ITRAP allows users to define which filtering steps to include, enabling to focus the filtering steps towards the generation of data with either high sensitivity or specificity. In summary, we believe ITRAP to be a highly valuable tool for the analysis of single-cell TCR specificity data, and we expect the tool to serve the community toward enabling an interpretation and denoising of such data resulting in an improved understanding of the underlying rules defining TCR specificity.

## Data Availability Statement

The different data sets analyzed and generated in this study are available at https://services.healthtech.dtu.dk/suppl/immunology/ITRAP_benchmark/. These data includes the raw data file (raw.cvs), the data filtered by the optimized UMI count thresholds (opt_thr.csv), the data filtered by the UMI thresholds and HLA matching (hla_match.csv), and final filtered data including only GEMs with complete TCR annotation (tcr.csv).

The data used for the ML training and evaluation of the two denoising pipelines is available at https://github.com/mnielLab/iTRAP_benchmark.

## Authors Contributions

M.N. conceived the idea. H.R.P and A.M. designed experiments, analyzed data, and made figures. L.E.J and M.N supervised the study. All authors wrote and revised the manuscript.

## Acknowledgements

The work was supported in part by National Institute of Allergy and Infectious Diseases (NIAID), under award number 75N93019C00001.

